# Different resource allocation in a *Bacillus subtilis* population displaying bimodal motility

**DOI:** 10.1101/2021.01.21.427716

**Authors:** Simon Syvertsson, Biwen Wang, Jojet Staal, Yongqiang Gao, Remco Kort, Leendert W. Hamoen

## Abstract

To cope with sudden changes in their environment, bacteria can use a bet-hedging strategy by dividing the population into cells with different properties. This so-called bimodal or bistable cellular differentiation is generally controlled by positive feedback regulation of transcriptional activators. Due to the continuous increase in cell volume, it is difficult for these activators to reach an activation threshold concentration when cells are growing exponentially. This is one reason why bimodal differentiation is primarily observed from the onset of the stationary phase when exponential growth ceases. An exception is the bimodal induction of motility in *Bacillus subtilis*, which occurs early during exponential growth. Several mechanisms have been put forward to explain this, including double negative-feedback regulation and the stability of the mRNA molecules involved. In this study, we used fluorescence-assisted cell sorting to compare the transcriptome of motile and non-motile cells and noted that expression of ribosomal genes is lower in motile cells. This was confirmed using an unstable GFP reporter fused to the strong ribosomal *rpsD* promoter. We propose that the reduction in ribosomal gene expression in motile cells is the result of a diversion of cellular resources to the synthesis of the chemotaxis and motility systems. In agreement, single-cell microscopic analysis showed that motile cells are slightly shorter than non-motile cells, an indication of slower growth. We speculate that this growth rate reduction can contribute to the bimodal induction of motility during exponential growth.

**IMPORTANCE:** To cope with sudden environmental changes, bacteria can use a bet-hedging strategy and generate different types of cells within a population, so called bimodal differentiation. For example, a *Bacillus subtilis* culture can contain both motile and non-motile cells. In this study we compared the gene expression between motile and non-motile cells. It appeared that motile cells express less ribosomes. To confirm this, we constructed a ribosomal promoter fusion that enabled us to measure expression of this promoter in individual cells. This reporter fusion confirmed our initial finding. The re-allocation of cellular resources from ribosome synthesis towards synthesis of the motility apparatus results in a reduction in growth. Interestingly, this growth reduction has been shown to stimulate bimodal differentiation.

## INTRODUCTION

Bacterial populations comprise different cell phenotypes, even when cells are genetically identical and exposed to the same environmental factors. For example, when the soil bacterium *Bacillus subtilis* is starved for nutrients, only a limited fraction of the population will become naturally transformable or will initiate sporulation. These developmental bifurcations are controlled by so called bimodal or bistable regulation pathways (1–3). Central to bimodal regulation is the presence of positive feedback control and a threshold level for activation of this positive feedback loop (4, 5). Random fluctuations in transcription and translation will lead in some cells to regulator concentrations reaching the threshold level, resulting in the stimulation of the positive feedback loop. In other cells the threshold level will never be reached. In this way two different cell types can develop in the same culture. The advantage of this heterogeneity is that a population as a whole can better adapt to unpredictable environmental changes. For example, when a starved and sporulating *B. subtilis* culture is suddenly exposed to a fresh influx of nutrients, sporulating cells cannot adjust to this since the sporulation program cannot be terminated (6). However, the non-sporulating cells are ready to use the new influx of nutrients to resume growth (7). Thus, during the process of sporulation, *B. subtilis* follows a bet-hedging strategy (1).

Bimodal regulation is wide spread. In fact, the induction of genetic competence, motility, biofilm matrix production and protease expression in *B. subtilis* are all controlled by positive feedback regulation cascades with a bimodal outcome (2, 3, 8–10). Many other bacteria employ bimodal regulation and for example pathogens like *Vibrio cholerae*, *Streptococcus pneumoniae*, *Salmonella typhimurium*, and *Pseudomonas aeruginosa* use it to create cooperative subpopulations for virulence (11–14).

The bimodal induction of sporulation, competence, matrix production and protease expression in *B. subtilis* occur when nutrients become limiting for optimal growth and cells enter the stationary growth phase. The bimodal induction of motility is curious since it occurs during exponential growth (8). Motility is also governed by positive feedback regulation and, in theory, the positive regulator should not easily reach an activation threshold concentration when cells are growing exponentially.

The motility system of *B. subtilis* is controlled by the sigma factor σ^D^, encoded by *sigD*. The *sigD* gene is located close to the 3’-end of the 26 kb long *fla/che* operon, encoding most of the flagellum and chemotaxis proteins. This operon is co-activated by two σ^D^-dependent promoters, thus providing the positive feedback necessary for bimodal induction (15). The position of *sigD* at the penultimate 3’-end of the very long *fla/che* transcripts makes expression of this gene prone to exonucleolytic mRNA degradation and premature transcription termination. Indeed, when *sigD* was placed close to the 5’-end of the operon, all cells activated the *fla/che* operon (15). It was therefore postulated that the stochastic fluctuation necessary for bimodality might be caused by fluctuations in RNA polymerase processivity and/or mRNA stability (15). Premature transcription of the SigD regulon is prevented by the SigD anti-sigma factor FlgM (16, 17), and the transcription factors SinR and SlrR that form a complex (18, 19). SinR alone represses *slrR* transcription and this activity is suppressed by binding of SlrR. It has been shown that this double negative feedback loop is also important for the bimodal induction of motility (20). Finally, SinR is also regulated by the protein SlrA that antagonizes binding of SinR to DNA (21, 22). Interestingly, the *slrA* transcript is directly targeted by polynucleotide phosphorylase (PNPase), the major 3’ exonuclease turnover enzyme, providing further support that RNA stability is a regulator of motility (23). It should be mentioned that the regulation cascade controlling motility is more complicated. For a comprehensive review see e.g. (24).

To obtain a possible clue as to why motility is induced during exponential growth, we separated *sigD* expressing cells from non-expressing cells using fluorescence-assisted cell sorting (FACS) and compared their transcriptomes. The data indicated that motile cells have a slightly but significantly reduced expression of ribosomal genes compared to non-motile cells. This difference was confirmed microscopically by the use of a reporter strain with an unstable variant of GFP. The reduction in ribosome synthesis pointed towards slower cell growth. Cell length measurements of motile and non-motile cells provided support for this conclusion, implying that synthesis of the motility system appears to exert a substantial pressure on the protein synthetic capacity of cells. We discuss how this growth reduction can contribute to bimodal induction during logarithmic growth.

## RESULTS

### Reciprocal distribution of motility and ribosomal genes

Exponentially growing *B. subtilis* cultures contain motile and non-motile cells (8). To determine whether these cells show any differences aside of the expression of SigD-activated motility and chemotaxis genes, we separated motile cells from non-motile cells by FACS and determined the transcriptome profiles of both fractions. In order to do this, we used reporter strain bSS339, containing the *gfp* gene under control of the strong SigD-activated flagellin (*hag*) promoter (8). It should be mentioned that this is a laboratory strain (168CA) that lacks the swarming motility capacity of undomesticated strains due to a mutation in *swrA* (25, 26). Cells were grown until an OD_600_ of ~0.5 after which the culture was subjected to cell sorting and separation into GFP-on (motile) and GFP-off (non-motile) cells (Fig. 1A-C). Motile cells induce autolysins involved in cell separation, which accounts for the differences in particle size (Fig. 1B) (27). The transcriptome profiles of both populations were determined by RNA-seq, and the experiment was repeated one more time. Table 1 lists the genes with significant (p-value <0.01) expression differences between motile and non-motile cells. A total of 22 genes were up-regulated and 33 down-regulated in motile cells compared to non-motile cells. 15 of the up-regulated genes are known SigD-activated genes (28). Interestingly, 15 of the down-regulated genes appeared to code for ribosomal proteins. When the expression differences of known ribosomal subunit genes and SigD-activated genes were plotted in a frequency histogram, a clear reciprocal distribution of motility genes and ribosomal genes is observed (Fig. 1 D). This was puzzling since the cultures used for FACS separation were growing exponentially with sufficient nutrients, and at a first glance there seems to be no reason for cells to lower their protein translation capacity.

**Fig. 1.**
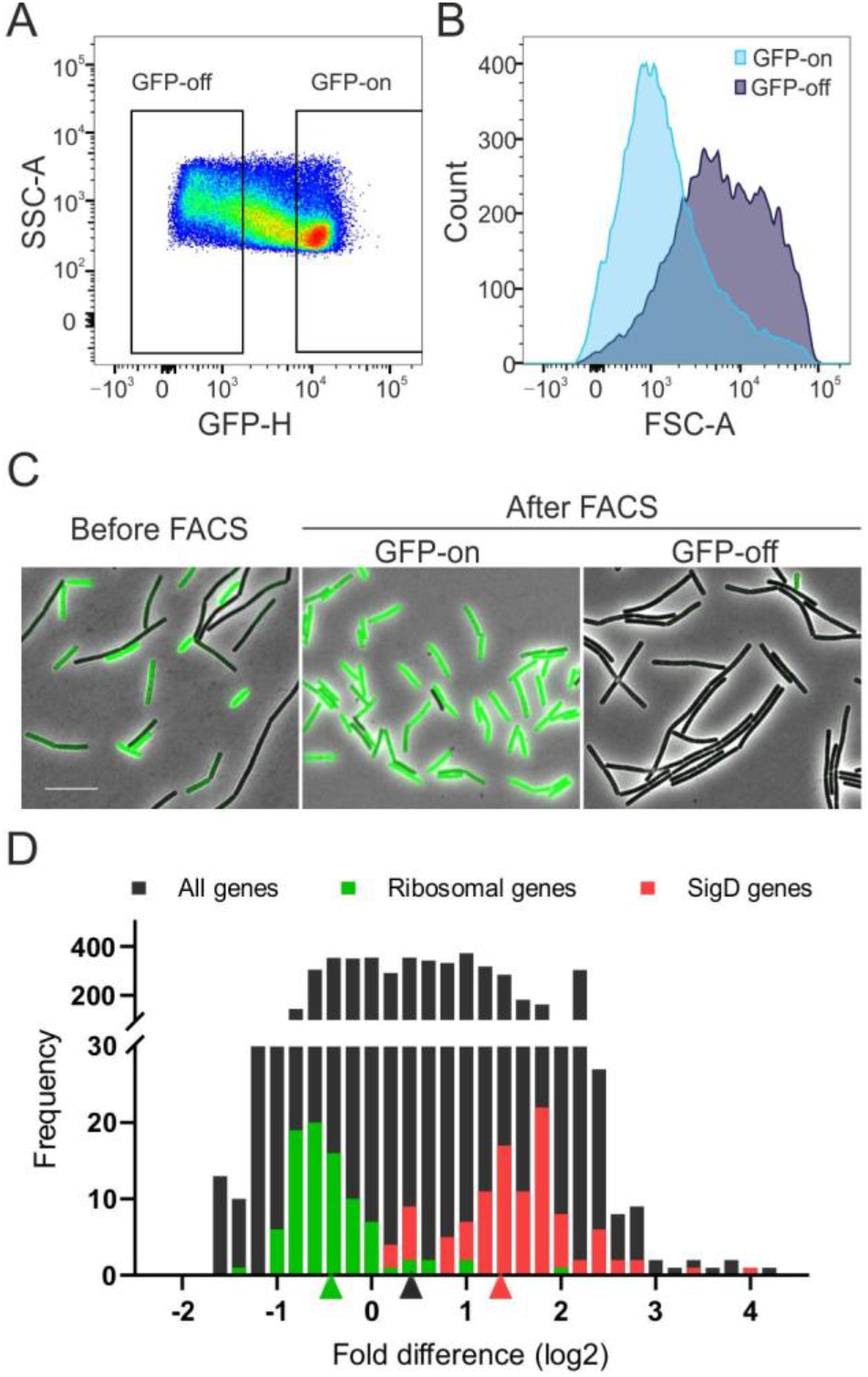
Fluorescence-activated cell sorting (FACS) for RNA-seq analysis of motile (GFP-on) and non-motile cells (GFP-off). Cells contained a GFP reporter gene under control of the flagellin promoter *Phag*. (A) Gating strategy (side scatter-area (SSC-A) versus fluorescence intensity (GFP-H) applied for flow cytometry sorting of GFP-on/off cells. (B) Size distribution of the GFP-on cells and the GFP-off cells based on forward scatter-area (FSC-A). (C) Microscopic images combining phase contrast and fluorescence of strain bSS339 (*Phag-gfp*) before and after FACS. (D) Histogram showing the distribution of ribosomal protein encoding genes (green) and SigD-activated genes (red) in the transcriptome profiles of motile cells compared to non-motile cells (GFP-on/GFP-off). All genes (black) were sorted on expression differences (X-axis), and binned according to measure of expression differences. 152 known SigD regulon genes and 53 ribosomal genes were used in the analysis (49). Arrows below the histogram show mean values of the three groups.

**Table 1.**
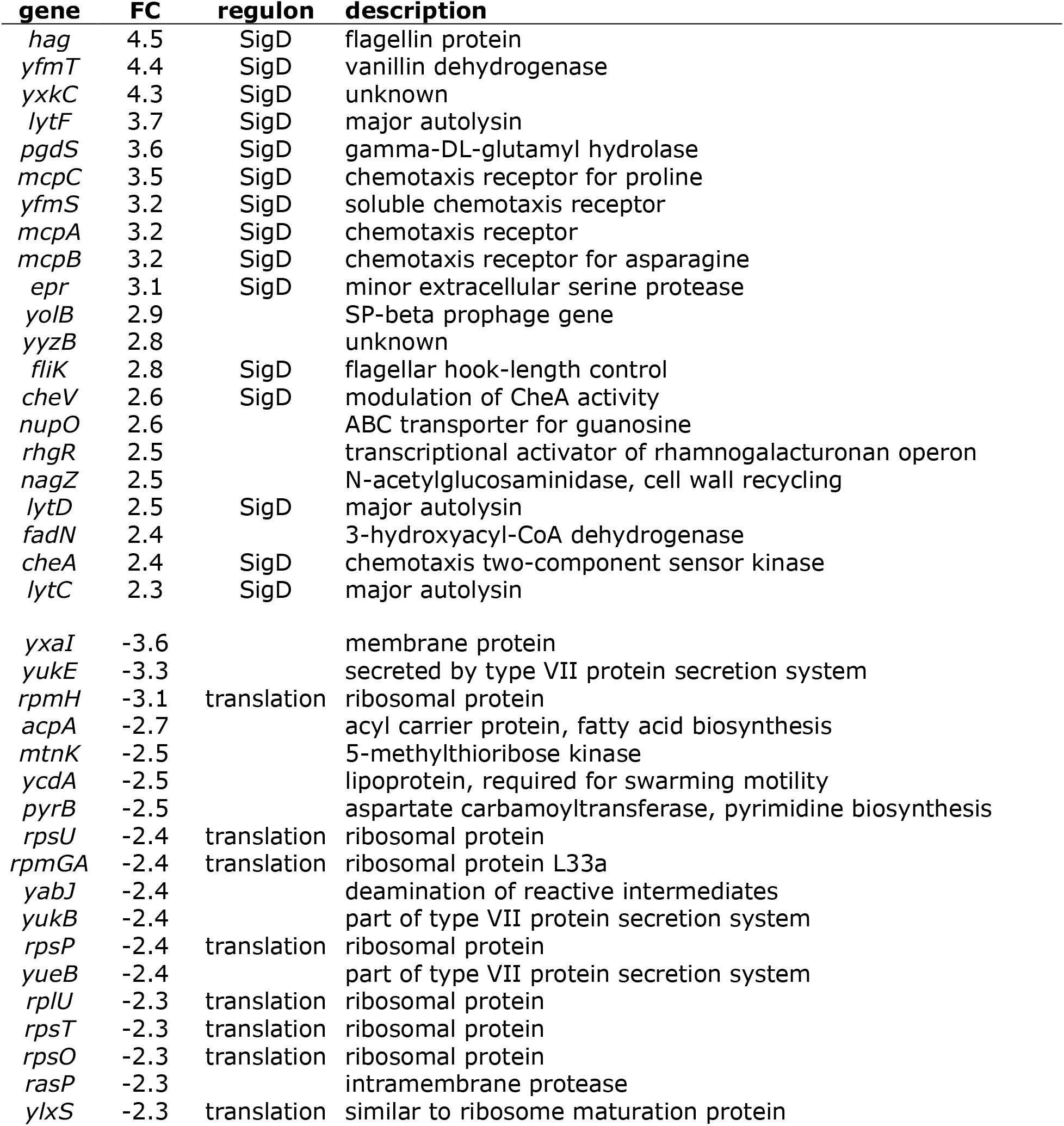
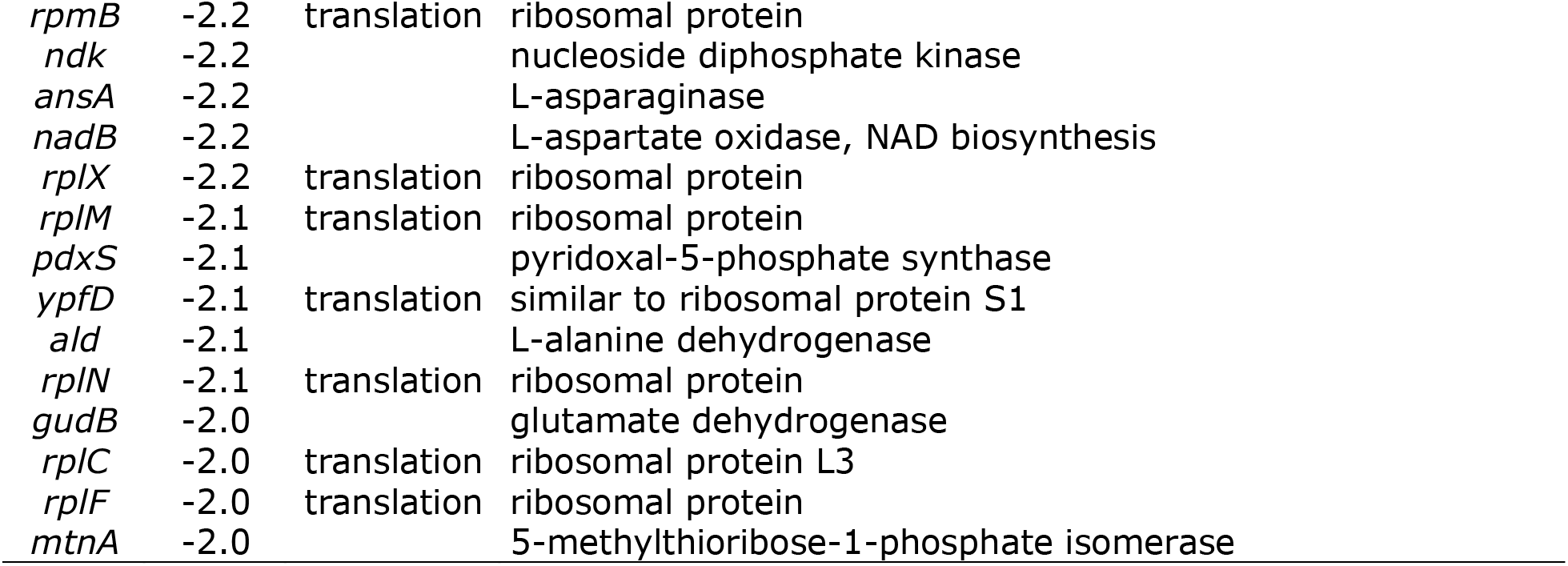
Transcriptome analysis of motile and non-motile cells. Genes are listed that showed a significant (p-value <0.01) fold change (FC) in expression between *hag* expressing (motile) and non-expressing (non-motile) cells based on two independent biological replicates.

### Instable GFP reporter

To confirm the reduced expression of ribosomal genes in motile cells, we monitored the activity of *rpsD*, coding for the ribosomal protein S4. A translational *rpsD* promoter (P*rpsD*) reporter fusion was constructed by fusing the first codon of the *rpsD* ORF in frame with *gfp*. The reporter fusion was constructed in an *amyE* integration vector, enabling ectopic expression. When the P*rpsD*-*GFP* reporter was introduced into *B. subtilis* all cells showed a very strong fluorescent signal without any obvious variation (data not shown). This is not surprising since the *rpsD* promoter is, like most ribosomal promoters, strongly expressed and GFP is a stable protein. Therefore, we created unstable variants of GFP by adding the *B. subtilis* SsrA tag (AGKTNSFNQNVALAA) to its C-terminus. Stalled proteins in the ribosome are cotranslationally tagged with a short peptide sequence encoded by the *ssrA* RNA before being released from the ribosome. This peptide tag targets the protein for degradation by the ClpXP protease complex (29). The addition of SsrA tag variants have been used before to destabilize GFP (30, 31). To test how SsrA variants affect GFP stability in *B. subtilis*, we used a *gfp* reporter expressed from the strong IPTG-inducible P*spac(*hy) promoter (32). When the SsrA peptide was fused to GFP, no fluorescent signal was detected (Fig. 2). Deleting *clpX* restored the fluorescence (Fig. 2). Clearly, the native SsrA tag is too efficient, therefore we tested two weaker SsrA tag variants in which the C-terminal 3 amino acids LAA were converted to AAV or DAV. These tags are less well-recognized by ClpX (30), resulting in intermediate GFP levels (Fig. 2).

**Fig. 2.**
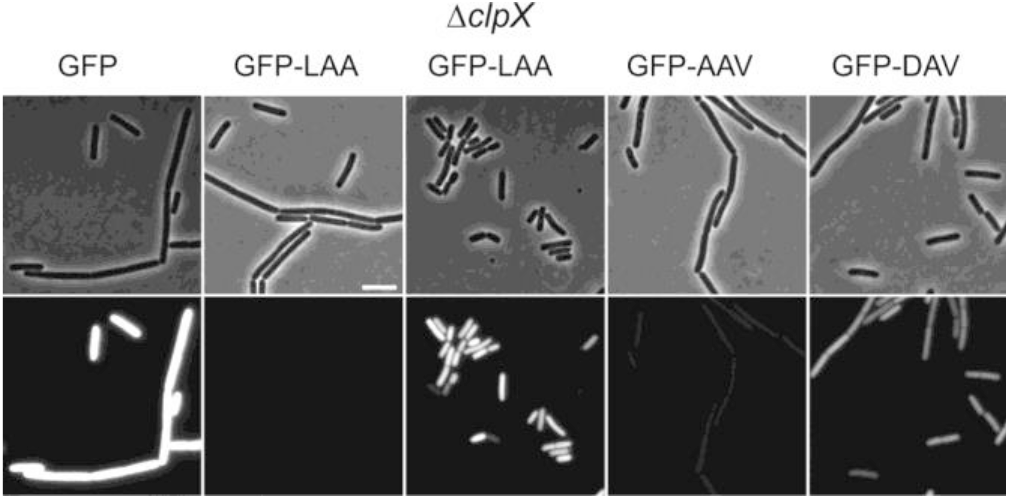
Effect of C-terminal SsrA tags on GFP stability. GFP was expressed from the strong IPTG-inducible P*spac(hy*) promoter induced with 1 mM IPTG. Fluorescence and related phase contrast images are shown in the bottom and top panels, respectively. The wild type SsrA tag ends with the sequence LAA. Strains used: bSS136 (GFP), bSS135 (GFP-LAA), bSS485 (GFP-LAA Δ*clpX*), bSS162 (GFP-AAV) and bSS161 (GFP-DAV).

Subsequently, the different SsrA tags were fused to the *PrpsD-GFP* reporter fusion. Interestingly, with this strong promoter, the wild type SsrA tag fusion still gave a fluorescence signal, albeit much lower compared to the other tags (Fig. 3). It should be mentioned that we have artificially increased the fluorescence signal intensity in Fig. 3 to improve the visibility, which explains the higher background signal for this strain. Apparently, the very strong expression from the ribosomal *rpsD* promoter is able to overcome the degradation rate by ClpXP. To measure the half-life of the different reporter fusions, translation was blocked by the addition of chloramphenicol, followed by microscopic measurements over time (Fig. 3, right panels). The half-life of untagged GFP was more than 2 hours. Addition of the DAV and AAV mutant tags reduced this to approximately 18 min and 10 min, respectively, and addition of the native SsrA tag reduced this further to less than 5 min.

**Fig. 3.**
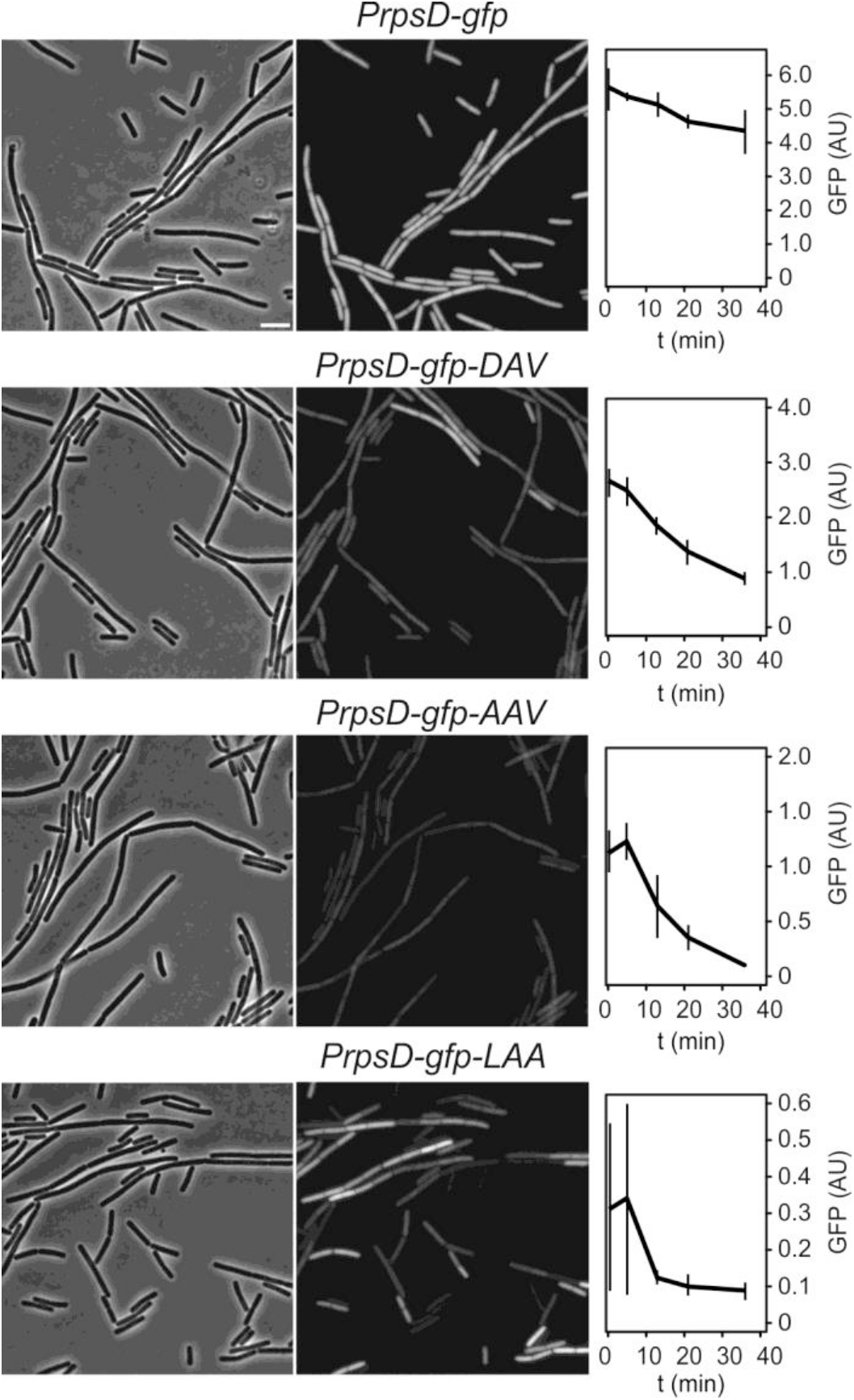
Half-life of GFP fused to different SsrA tags and expressed from the strong ribosomal *rpsD* promoter. Chloramphenicol was added at t = 0 min. The calculated half-life for GFP without an SsrA tag, and with either the SsrA-DAV, SsrA-AAV or wild type SsrA-LAA tag, is 120 min, 18 min, 10 min and 5 min, respectively. The wild type SsrA tag ends with the sequence LAA. Strains used bSS429 (*PrpsD-gfp*), bSS492 (*PrpsD-gfp-DAV*), bSS432 (*PrpsD-gfp-AAV*) and bSS387 (*PrpsD-gfp-LAA*).

### Reduced *rpsD* expression in motile cells

As shown in Fig. 3, the *B. subtilis* strain harboring the GFP variant with the shortest half-life displayed the largest heterogenic expression in cells, and this reporter strain was further used to monitor differences in ribosome synthesis between cells. To examine whether motile cells have a reduced expression of ribosomal genes, as the transcriptome data suggested, the P*rpsD-GFP-ssrA* reporter (green fluorescence), was introduced into a strain containing a red fluorescent reporter protein, mCherry, expressed from the SigD-activated flagellin promoter (*Phag*-*mCherry*). Fig. 4B shows a composite image of the green and red fluorescence channel from a sample taken early during exponential growth (OD_600_ ~0.2). Non-motile cells, often arranged in long cell chains, show on average higher GFP signals compared to the red fluorescent motile cells (Fig. 4B, arrow heads). The cell chaining is caused by a low levels of the autolysin LytF, which is expressed when SigD is active (27). Fig. 4C-F show scatter histograms of the red and green fluorescence signals of cells sampled during growth. From the distribution of data points it is apparent that there is a reciprocal relation between the activity of the *rpsD* promoter and the *hag* promoter during early exponential growth (time points C, D and E). To clarify this, cells were subdivided into a SigD-on and -off population on the X-axis using the background red fluorescence of a Δ*sigD* mutant as cutoff. Division on the Y-axis into high and low *rpsD* activity was based on the background GFP fluorescence of the stationary phase culture, which showed low P*rpsD* activity (Fig. 4C-E, cell numbers in right upper corner). Based on this division more than 85 % of cells with a high *rpsD* activity fall within the SigD-off population, during exponential growth. This distribution disappears in stationary phase (Fig. 4F) since the expression of ribosomal genes, including *rpsD* is strongly reduced when nutrients become limiting for growth.

**Fig. 4.**
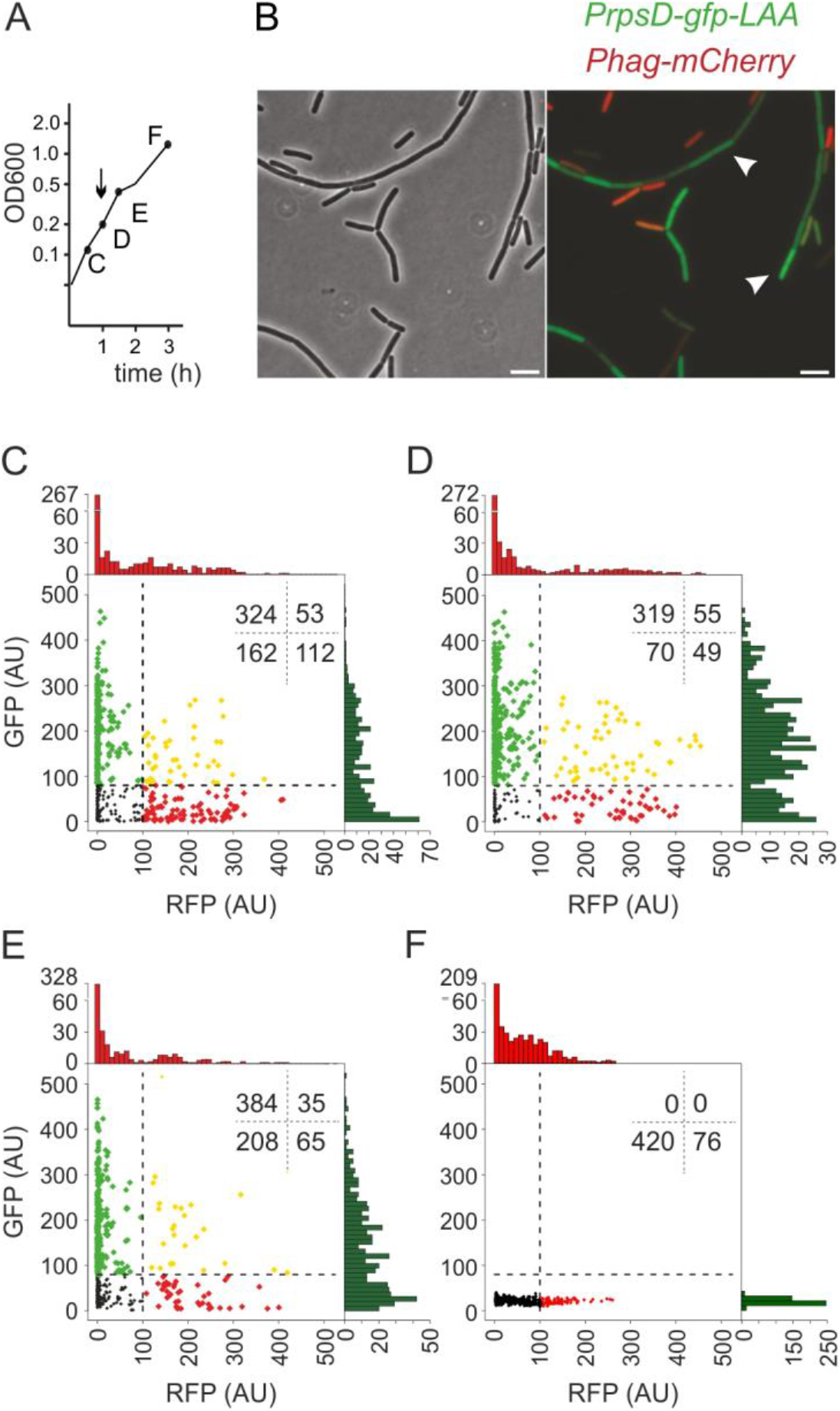
Reciprocal relation between SigD and ribosomal protein RpsD expression during growth. SigD expression is monitored by following the expression of the SigD-dependent *hag* promoter fused to mCherry. RpsD expression is monotored by fusion of the *rpsD* promoter to GFP tagged with LAA, which reduces the half-life of GFP drastically. (A) Cells were grown in CH medium at 37 °C and samples for mycroscopic analysis were taken after 30, 60, 90, and 180 min. (B) Phase contrast and merged fluoresence image showing the mCherry and GFP signals. The image was captured after 60 min. White arrow heads indicate two non-motile cell chains. Scale bar is 5 μm. (C–F) Scatter-histogram plots of P*rpsD-gfp-ssrA* and P*hag-mCherry* activities of individual cells from the four time points indicated in panel A. Cells are subdivided into P*hag*-on and -off populations on the x-axis, and P*rpsD*-on and off populations on the y-axis. See main text for the determination of threshold values for on and off populations for the two reporters. Numbers in the top right corners signify cell counts in the four quadrants. Strain used bSS387 (P*rpsD-gfp-ssrA* P*hag-mCherry*).

The expression of *rpsD* is auto-regulated by binding of its gene product, ribosomal protein S4, to the leader region of its own mRNA (33, 34). Inactivation of this autoregulation by removing the S4 binding region from *rpsD* did not change the results (Fig. S1), suggesting that the reciprocal relation between *rpsD* expression and motility induction occurs primarily at the level of *rpsD* transcription.

As a final test, we examined whether inactivation of SigD affects the expression of ribosomal genes during exponential growth. Using q-RT-PCR, the expression levels of *rpsD* and two other ribosomal genes, *rpsJ* and *rplC* encoding ribosomal protein S10 and L3, respectively, were monitored in wild type and Δ*sigD* cells (Fig. S2). Indeed, the mRNA levels of these three genes were 2 to 6 times lower in a wild type culture compared to a Δ*sigD* culture. These data confirmed the transcriptome data, indicating that cells reduce their protein translation capacity when they become motile, even when nutrients are plentiful.

### Growth rate reduction

The lower expression of ribosomal genes in motile cells suggested that these cells are growing slower compared to non-motile cells. Growth rate is directly related to cell size (35). Indeed, when we measured the cell length of motile and non-motile cells, using the red fluorescent membrane dye Nile red to distinguish individual cells in chains of *B. subtilis* cells (36), a clear difference in cell length was observed, with motile cells being on average 15 % shorter than non-motile cells (Fig. 5A, B).

**Fig. 5.**
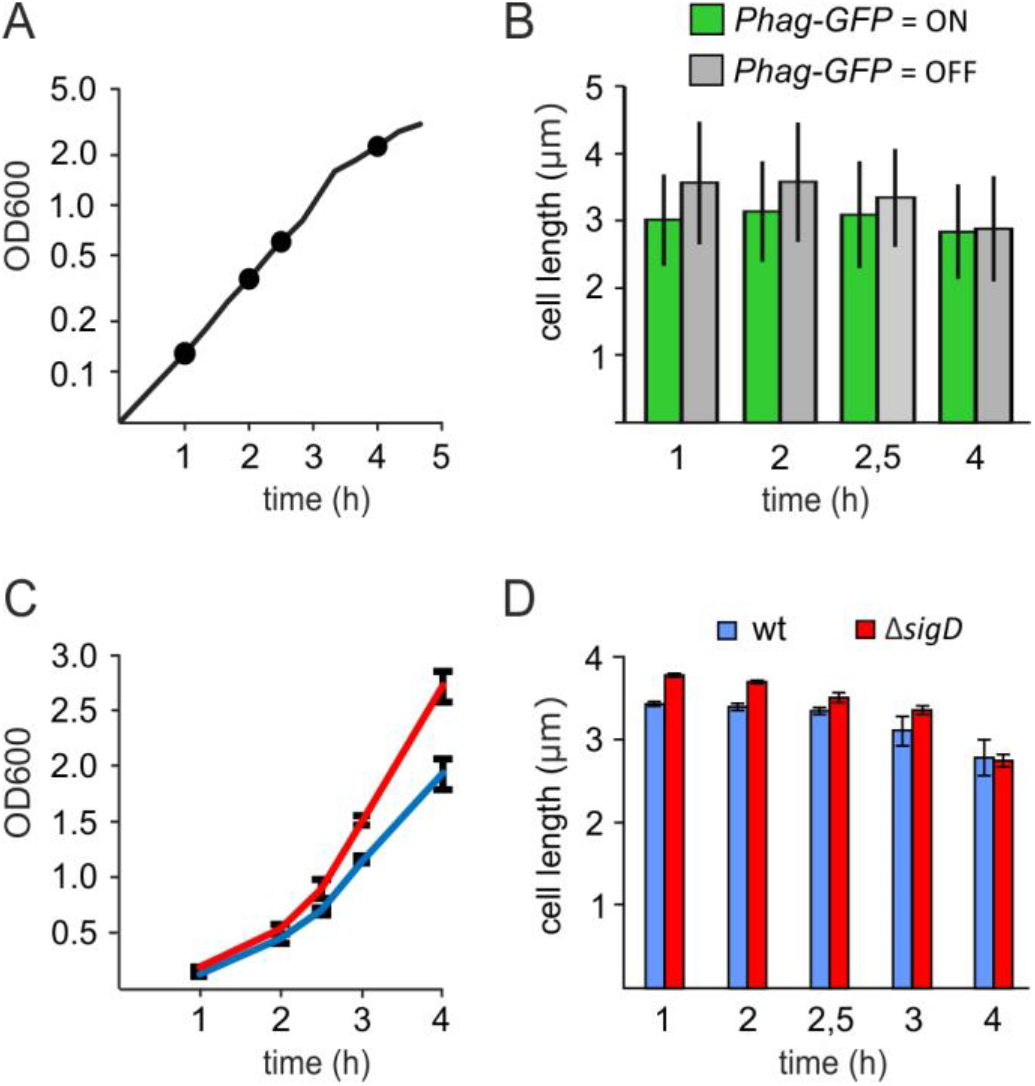
Effect of SigD induction on cell length. (A) Time points for cell length measurements of cells grown in CH medium at 37 °C. (B) Average cell length of SigD-on and -off cells. SigD activity was determined by P*hag-GFP* expression, and the threshold value for on/off determination was 100 AU GFP based on data from Fig. 4. Values were based on measurement of approximately 400-500 cells per time point. (C) Average growth curves of wild type and Δ*sigD* mutant in CH medium at 37 °C indicating the time point for cell length measurements. (D) Average cell length of wild type and Δ*sigD* mutant cells measured at time points indicated in (C). Values were based on three biological replicates and measurements of approximately ~500 cells per time point. Strains used bSS339 (*Phag-gfp*), 168 (wild type) and CB100 (Δ*sigD*).

The simplest explanation for this is the impact the synthesis of the motility and chemotactic sensing system has on cell physiology. SigD activates expression of at least 151 proteins (28, 37), among which the flagellin protein Hag, which is one of the most abundant proteins in *B. subtilis* (38). As a result, cells will use fewer resources to express constitutive genes, including those responsible for cell growth. Therefore, the SigD-off cells will be longer compared to SigD-on cells, assuming that the rate of cell division is constant. If this is true, then blocking induction of motility should increase the growth rate. To test this, we compared the growth rate of a Δ*sigD* mutant with a wild type strain (Fig. 5C). Although the differences are small, a Δ*sigD* mutant grows slightly faster and cells are a bit longer (Fig. 5D), confirming that the induction of chemotaxis and motility puts a burden on the cell’s transcriptional and translation resources.

## DISCUSSION

In this study we show that the induction of motility in *B. subtilis* puts a strain on available cellular resources, diverting transcription and translation capacity from synthesis of ribosomes to synthesis of the motility and chemotaxis systems. As a result of this, non-motile cells grow faster than motile cells. Although this effect has been predicted (e.g. (39)), to our knowledge, this is the first time that this phenomenon has been directly measured.

We could only observe expression effects on the ribosomal *rpsD* promoter when the GFP reporter was made unstable by fusing it to the SsrA degradation tag. Therefore, we cannot exclude with certainty that the variability in cellular expression is not related to cellular variations in the activity of the protease ClpXP complex. However, if this would be the case, it seems likely that the other, less active, SsrA tag variants also would show some measure of heterogeneity, and this was not detected.

Normally, exponential growth of cells will reduce the chance of a positive feedback loop to be activated due to the dilution of the activator. In fact, bimodal expression occurs generally under suboptimal growth conditions, such as in the stationary growth phase. In *B. subtilis* this is for example the case for the development of genetic competence, sporulation, and secretion of proteases (1–3, 10). However, the bimodal induction of *sigD* activation during exponential growth might actually be related to its effect on growth rate. During balanced growth there is an inverse correlation between growth rate and the concentration of constitutively expressed proteins in the cell, whereby higher growth rates result in lower protein concentrations in the cell (40). The reason for this is that the volume of a cell increases exponentially faster than the copy number of constitutively expressed genes in the cell (DNA replication), including those of ribosome encoding genes. This inverse relation between growth rate and protein expression can create a positive feedback mechanism for the expression of proteins that affect growth, since a reduction of the growth rate will increase protein concentration and translation capacity. This can lead to bimodal expression and related bimodal growth (40). For example, when in *E. coli* the strong T7 polymerase was expressed from a T7 promoter, this simple positive feedback loop resulted in a bimodal response with only a subset of *E.coli* cells producing T7 polymerase during exponential growth (41). Presumably, something similar occurs with the activation of *sigD* in *B. subtilis* during exponential growth. Due to leaky expression, SigD levels can be slightly higher in some cells compared to others. In these cells, the stimulated expression of SigD activates at least 150 genes, which will divert transcription and translation capacity away from the expression of growth related proteins, resulting in a reduction of the growth rate. This will be further enhanced due to the competition between the housekeeping sigma factor (SigA) and SigD for RNA polymerase holoenzyme. Subsequently, this reduction will increase the concentration of proteins, including SigD, which will then further stimulate the expression of SigD-regulated genes, among which its own gene, resulting in bimodal expression during exponential growth.

Bimodal induction of *sigD* is the result of many factors that influence expression of *sigD*. Aside from the possible growth rate reduction-based feedback shown here, there is also the activity of the SigD anti-sigma FlgM. Cellular FlgM levels are reduced due to secretion of the protein through the flagellum basal body, and this in itself presents a positive feedback loop (15). As mentioned in the introduction, there is also the double negative regulatory feedback loop composed of SinR and SlrR, and the mRNA stability of the long 27 kb *fla*/*che* operon that affects expression of *sigD*. The possibility to spread out and reach different niches requires motile cells to be more adaptive than sessile cell chains that build up biofilms, which may explain the complex regulation, comprising several bimodal feedback systems.

## MATERIALS AND METHODS

### Strains and growth conditions

Strains were grown overnight in casein hydrolysate (CH) medium (42) containing 1 % glucose to inhibit sporulation, and then diluted 100x into fresh CH medium until OD_600_ 0.5. Samples were diluted again into pre-warmed fresh CH medium to a final OD_600_ of 0.05 and grown at 37°C. Samples were taken at indicated time points for OD_600_ measurements, FACS and microscopy analysis.

### Strain construction

Molecular cloning, PCRs, and transformations were carried out by standard techniques. Details of the strain construction are described in the supplementary information. Strains, plasmids and oligonucleotides are listed in Table S1, S2 and S3, respectively.

### FACS conditions

Strain bSS339 containing the P*hag-gfp* reporter fusion was used to separate motile (GFP-on) and non-motile (GFP-off) cells. Cells were cultured in 10 ml CH medium in 125 ml flasks to an OD_660_ of 0.5 and quickly cooled by adding 10 ml ice-cold PBS with 4 M NaCl, which preserves mRNA (43), and placed on ice. The culture was directly sorted by BD FACSAria^TM^ III flow cytometer with a 70 Micron Nozzle. During the sorting process the sample input and collection chamber were cooled and kept at 4°C. GFP-on/off cells were sorted using a 4-way purity precision mode, but mainly based on GFP intensity in the SSC-A scatter plot. The parental wild type *B. subtilis* 168 strain was used as GFP negative control. The GFP-on/off cells were sorted in 1X PBS flow stream and collected using 15 ml tubes half-filled with 4 M NaCl 1X PBS (44). The collected cells were mixed with the same volume of 4 M NaCl 1X PBS. The tubes were then centrifuged for 30 min at 4 °C at 20,000 rpm and the cell pellets were washed once with ice-cold PBS, flash-frozen with liquid nitrogen and stored at −80 °C for further processing. The flow-cytometry data were analyzed using FlowJo v10 software (FlowJo LLC, Ashland, OR). Subsequent RNA isolation and RNA-seq analyses are described in the supplementary information. RNA-seq data have been submitted to and are accessible in the Gene Expression Omnibus (GEO), accession number GSE164348

### Microscopy

Cells were stained with the fluorescent membrane dye Nile Red (0.2 μg/ml) and spotted on a thin 1 % agarose slab. A Zeiss M200 microscope using a CoolSNAP HQ2 CCD camera and Metamorph v.7.7.5.0 was used to capture images. Exposure times for GFP and Nile red were 500 msec, and 250 msec, respectively. Images were analyzed using ImageJ (National Institutes of Health) and R (R foundation for statistical computing). The Image J Plugin ChainTracer was used to measure cell length (36).

### q-RT-PCR

RNA isolation for quantitative reverse transcription PCR was performed based on (45) and (46). Briefly, cells were collected at 1 and 2 h into exponential growth, corresponding to OD_600_ 0.2 and 0.6, respectively. RNA was extracted as described in the supplementary information. RNA was reverse-transcribed into cDNA using Thermo Fisher’s First Strand cDNA synthesis kit. qPCR was executed using Thermo Fisher DyNAmo HS SYBR-Green qPCR kit and an Applied Biosystems 7500 Real-Time PCR system. bSS387 and bSS407 were used as wild type and Δ*sigD* strain, respectively. Primers used are listed in Table S3. Relative quantification of ribosomal genes was performed using the 2^-ΔΔCT^ method, taking *gapA*, coding for glyceraldehyde 3-phosphate dehydrogenase, as an internal control (47, 48).

## Acknowledgements

We would like to acknowledge Wayne Rasband for developing and maintaining ImageJ. We thank Daniel Kearns for strains. SS was supported by a BBSRC DTG PhD studentship 2009, YG and BW were supported by a China Scholarship Council fellowship, and LWH was supported by an NWO STW-Vici (12128) grant.

